# MCE domain proteins: conserved inner membrane lipid-binding proteins required for outer membrane homeostasis

**DOI:** 10.1101/159053

**Authors:** Georgia L. Isom, Nathaniel J. Davies, Zhi-Soon Chong, Jack A. Bryant, Mohammed Jamshad, Maria Sharif, Adam F. Cunningham, Timothy J. Knowles, Shu-Sin Chng, Jeffrey A. Cole, Ian R. Henderson

## Abstract

Bacterial proteins with MCE domains were first described as being important for <u>M</u>ammalian <u>C</u>ell <u>E</u>ntry. More recent evidence suggests they are components of lipid ABC transporters. In *Escherichia coli*, the single-domain protein MlaD is known to be part of an inner membrane transporter that is important for maintenance of outer membrane lipid asymmetry. Here we describe two multi MCE domain-containing proteins in *Escherichia coli*, PqiB and YebT, the latter of which is an orthologue of MAM-7 that was previously reported to be an outer membrane protein. We show that all three MCE domain-containing proteins localise to the inner membrane. Bioinformatic analyses revealed that MCE domains are widely distributed across bacterial phyla but multi MCE domain-containing proteins evolved in Proteobacteria from single-domain proteins. Mutants defective in *mlaD*, *pqiAB* and *yebST* were shown to have distinct but partially overlapping phenotypes, but the primary functions of PqiB and YebT differ from MlaD. Complementing our previous findings that all three proteins bind phospholipids, results presented here indicate that multi-domain proteins evolved in Proteobacteria for specific functions in maintaining cell envelope homeostasis.

---

Mammalian cell entry (MCE) domains are conserved amino acid motifs that are widespread across bacteria^1^. They were first identified when a chromosomal fragment containing the *mce1A* gene from *Mycobacterium tuberculosis* was inserted into non pathogenic *Escherichia coli,* which allowed the latter bacterium to enter and survive in mammalian cells^2^.

Research on MCE domain-containing proteins (henceforth termed ‘MCE proteins’) has largely focused on elucidating their roles in Actinobacteria. *In silico* analysis indicated that the *mce* operons in Actinobacteria encode ABC transporters^3^ as they often encode permease domains and occasionally ATPase domains^4^, which are typical components of an ABC transporter^5^. Similarities have been demonstrated between MCE domains and the substrate binding proteins of ABC transporters^4^. A more specific role in lipid transport has been suggested, as evidenced by research linking *mce* operons in *M. tuberculosis* to the transport/uptake of lipids^6-11^. All Actinobacterial MCE proteins studied thus far contain a single MCE domain.

MCE domains in Proteobacteria have recently received considerable attention, with particular focus on the single MCE domain protein MlaD, which forms part of an ABC transporter complex found in the inner membrane^12,13,14^. In *E. coli*, this Mla pathway has been shown to play a role in outer membrane maintenance through trafficking of phospholipids from the cell surface back into the cell^12^. The outer membrane of Gram-negative bacteria is asymmetric, with lipopolysaccharides (LPS) and phospholipids found in the outer and inner leaflets, respectively. The Mla pathway in *E. coli* is important for maintaining this lipid asymmetry as mutations in the pathway, including MlaD, result in phospholipid accumulation in the outer leaflet of the outer membrane^12^. This defect is elevated in cells lacking *pldA*, which encodes an outer membrane phospholipase that degrades phospholipids in the outer leaflet^12^. Similar pathways have been studied in other Proteobacteria^15-17^ and also in the plant species *Arabidopsis thaliana*, where the ABC transporter traffics phosphatidic acid in the chloroplast^18^.

Two other proteins in *E. coli*, PqiB and YebT, are predicted to contain MCE domains^19^. Unlike Actinobacterial MCE proteins, they contain multiple MCE domains (3 and 7, respectively) and their functions are currently unknown. The best-studied multi-MCE domain-containing protein is multivalent adhesion molecule 7 (MAM-7) in *Vibrio parahaemolyticus*, which is an orthologue of YebT. MAM-7 is reported to be an integral outer membrane protein on the cell surface that acts as an adhesin by binding to mammalian cells via phosphatidic acid and fibronectin^20^.

Given the wide distribution of MCE domains across Proteobacteria, we sought to understand functional and evolutionary relationships between these proteins, with a particular focus on those found in *E. coli.* Our analysis suggests that multi-domain proteins evolved from single domain proteins within Proteobacteria, potentially as part of a novel type of transporter located in the inner membrane. The data presented below, together with our recent findings that all three proteins bind phospholipids^13,14^, provides evidence that PqiB and YebT are involved in the transport of phospholipids and maintenance of outer membrane asymmetry, yet have ultimately evolved a primary function that differs to that of MlaD.

## Results

### MCE protein architectures and phylogenetic prevalence

Lipid shuttling between the inner and outer membranes of diderm bacteria is a poorly understood process. One protein known to be involved in this process is the *E. coli* protein MlaD. MlaD contains a single MCE domain; the PFAM hidden Markov model (HMM) for the MCE domain (PF02470.15) defines an 81 amino-acid long sequence with multiple well-conserved hydrophobic residues^1^. We hypothesised that other proteins involved in lipid shuttling might contain similar MCE domains. Thus, we sought to determine the distribution of these domains amongst bacteria and to investigate the most common MCE protein architectures by scanning the PFAM HMM against the UniProtKB database using HMMER^21^. A total of 17,282 MCE proteins were identified and scanned for other PFAM domains to construct protein architectures. Four architectures account for >97% of identified MCE proteins. They are found in 24 of the 31 phyla analysed and all contain an N-terminal transmembrane helix consistent with proteins known to localise to the cytoplasmic membrane (Fig. 1). These phyla primarily consist of diderm bacteria, including the Negativicutes, a subset of Gram-negative organisms found within the phylum Firmicutes. A full list of architectures can be found in Supplementary table S1.

**Figure 1.**
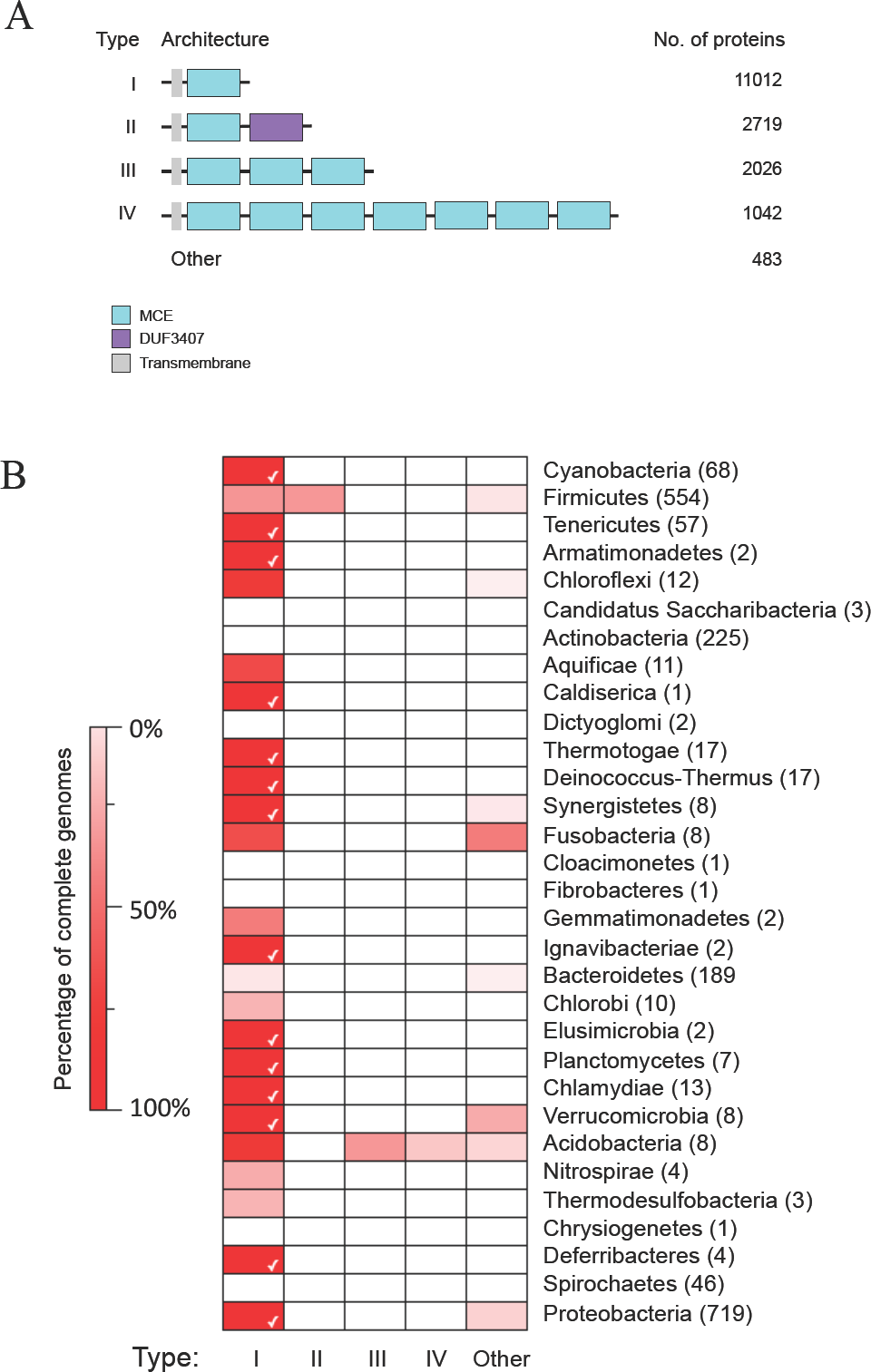
The distribution of the top MCE protein architecture across bacteria. (A) The top four MCE protein architectures by number of proteins. These were identified by scanning the UniprotKB protein database with the MCE hidden Markov model and other models from PFAM. (B) A heat map showing the distribution of the architectures across bacterial phyla, to include all species with a full genome sequence. The colours are based on percentages ranging from 0% (white) to 100% (bright red). The phyla are displayed on the right of the heat map and the number of species is in brackets. The phyla are ordered based on the tree of life^32^, ranging from early (top) to late (bottom) branching bacteria. A white tick is displayed when an architecture is found in 100% of the analysed species within a phylum.

Type I MCE proteins containing a single MCE domain and no other predicted PFAM domain are by far the most abundant and widely distributed. This group includes well-known proteins such as *E. coli* MlaD and multiple predicted lipid transport proteins in Actinobacteria such as Mce4B from the predicted cholesterol uptake pathway in *M. tuberculosis^9^.* The second most common architecture, type II, consists of a single MCE domain followed by a DUF3407 (Cholesterol Uptake Porter) domain, and is specific to Actinobacteria. DUF3407 domains are also specific to Actinobacteria and are almost always (>99%) associated with a single MCE domain^1^. Type III and IV proteins contain three and seven MCE domains in tandem, respectively. Both types are specific to Proteobacteria; in *E. coli* these have been designated PqiB (type III) and YebT (type IV). Type III proteins are more prevalent than type IV, which are restricted to Deltaproteobacteria, Epsilonproteobacteria, and Gammaproteobacteria (Supplementary fig. S1). Other multi-MCE domain-containing proteins were detected in Proteobacteria, but at much lower frequencies than type III and IV proteins (Supplementary table S1, Supplementary fig. S1).

Some MCE domains were detected in proteins from eukaryotic genomes (Supplementary fig. S2). Type I proteins were found in plant phyla Chlorophyta and Streptophyta. In *Arabidopsis* they are involved in the trafficking of phosphatidic acid from the outer to the inner membrane of chloroplasts^18^. A small number of MCE proteins were identified in animal genomes. Manual inspection of the DNA sequences encoding these proteins revealed that all but one could be attributed to contamination with bacterial DNA, the exception being in *Trichoplax adhaerens*, an animal known to have an unusually large mitochondrial genome^22,23^.

### Protein clustering and evolution of multi-domain proteins

To understand the evolutionary relationships between MCE proteins, protein-protein similarity networks were constructed and coloured by architecture type and phylum (Fig. 2). MCE proteins generally cluster within phyla, suggesting that little or no horizontal transmission of these genes has occurred and variants have arisen through speciation. Actinobacterial MCE proteins, whether type I or II, form a single tight cluster, perhaps suggesting functional homogeneity and purifying selective pressure in this phylum. Type I proteins from most other phyla cluster loosely, including Cyanobacteria, Bacteroidetes and a subset of Proteobacterial proteins. All plant MCE proteins cluster closely with Cyanobacteria, and the single *Trichoplax* protein with Proteobacteria, which is expected given the origins of chloroplasts^24^ and mitochondria^25^, respectively. The position of types III and IV proteins in the large Proteobacteria-dominant cluster suggests that these proteins are a functionally divergent population that arose early in Proteobacterial evolution.

**Figure 2.**
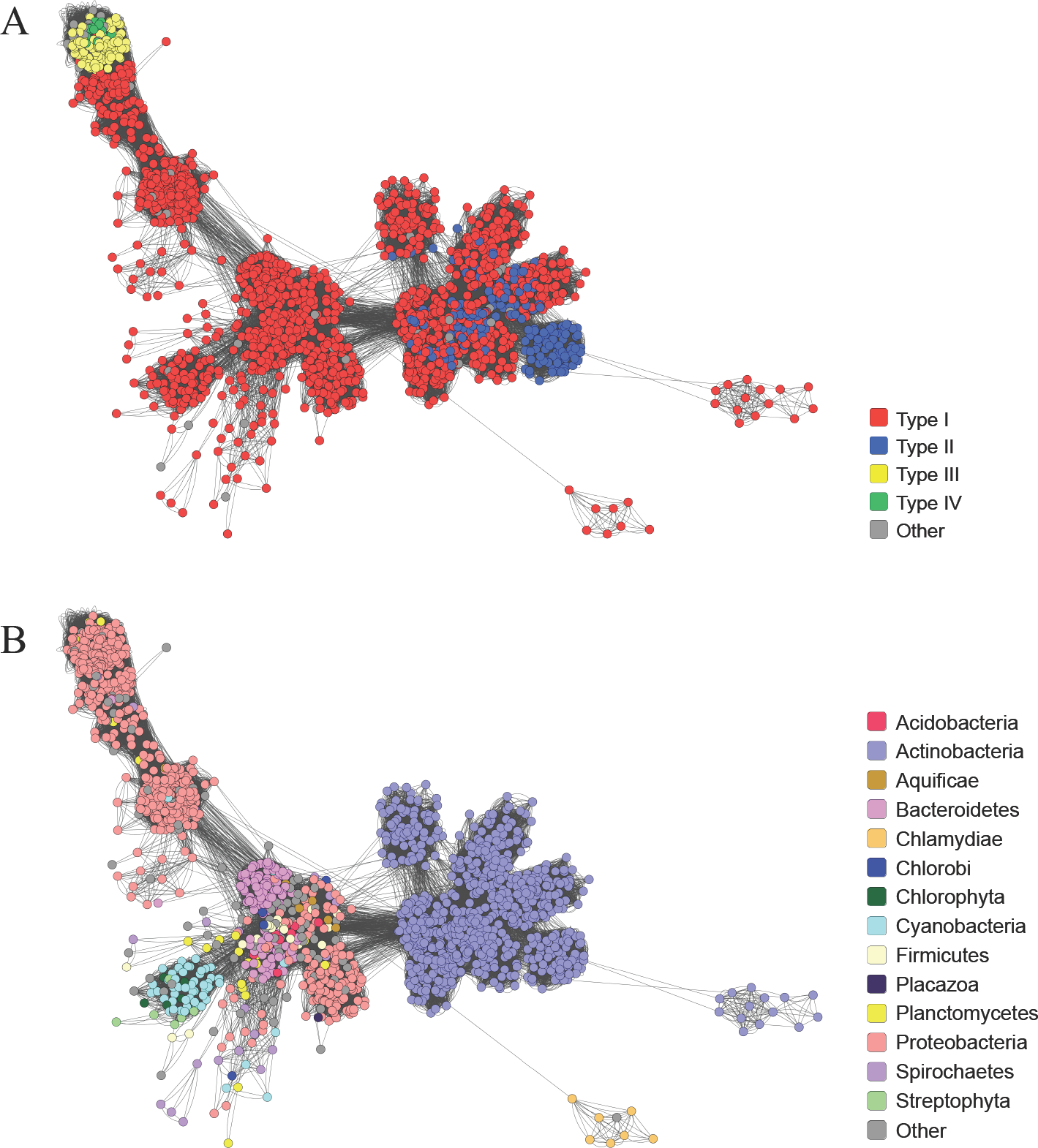
Sequence similarity clustering of all MCE proteins. A representative subset of MCE proteins (including those found in eukaryotes) were clustered based on similarity, as determined by BlastP. A node represents a protein and an edge (line) indicates an interaction between two proteins. The cluster diagram is coloured by (A) protein architecture and (B) phylum.

### Genes encoding MCE proteins co-localise with genes encoding membrane transport proteins

Based on our previous structural data we hypothesised MCE proteins formed part of a supramolecular complex spanning the cell envelope^14^. To identify proteins that might be functionally linked to MCE domains, the gene neighbourhoods of type I, III and IV genes in Proteobacteria were analysed independently (Fig. 3A). Type I genes are commonly (59%) found with permease and ATP-binding domains upstream. The most common downstream domain is the ABC auxiliary lipoprotein domain, DUF330. This domain is primarily found in Proteobacteria, and can exist as a gene fusion with genes encoding MCE domains (Supplementary table S1). In Beta- and Gammaproteobacteria, the DUF330 gene may be absent from the operon and in these cases genes encoding the periplasmic Tol domain and the cytoplasmic STAS domain are frequently located immediately downstream of the MCE protein encoding gene. The former domain is related to toluene resistance^16^ whilst the latter is a general NTP binding domain^26^. This operon type encodes the Mla pathway in *E. coli*^12^ (Fig. 3B). In many of these neighbourhoods, for example in *Neisseria meningitidis*^27^, the outer membrane component MlaA (VacJ) is found in the same operon.

**Figure 3.**
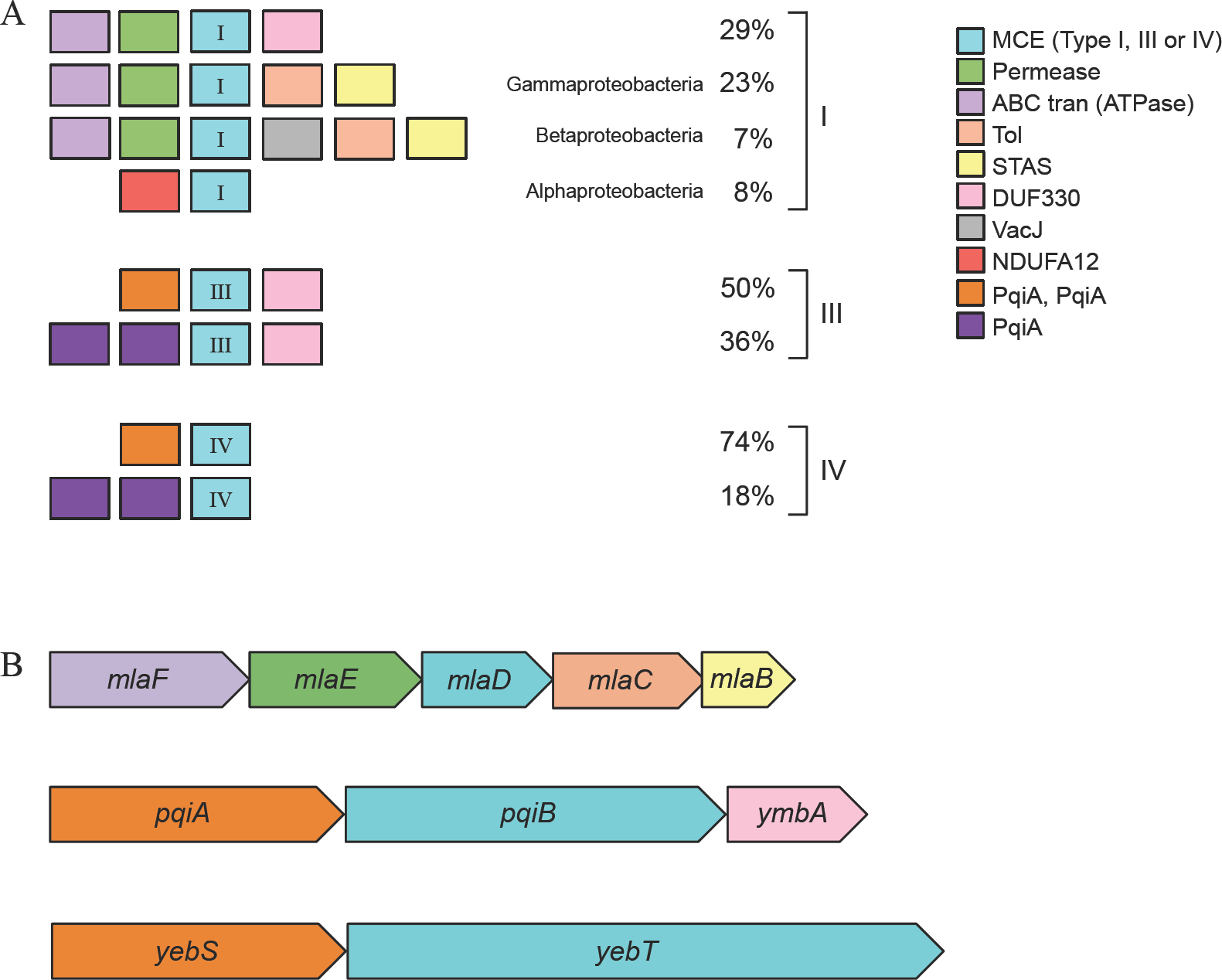
The predicted gene neighbourhoods of MCE-domain encoding genes in Proteobacteria. (A) The most common neighbourhoods in Proteobacteria separated by protein architecture. The percentage represents the occurrence of each neighbourhood for a given architecture. To identify the neighbourhoods, the 10 kb regions up and downstream of the MCE-domain encoding genes were translated and scanned for all PFAM domains. (B) The operons that encode type I (MlaD), type III (PqiB) and type IV (YebT) protein in *E. coli*.

In Alphaproteobacteria, an NADH dehydrogenase subunit domain (NDUFA12) is found upstream of a type I gene instead of ABC transporter proteins. Although it is unknown whether these genes are functionally related in bacteria, the same two domains are found together in the *T. adhaerens* MCE protein, suggesting that a gene fusion event might have occurred following the formation of mitochondria from Alphaproteobacteria^25^.

Like many type I proteins, but different to type IV proteins, type III proteins are commonly associated with genes encoding DUF330 domains, highlighting a distinction between the multi-domain proteins (Fig. 3A). The majority of the type III and type IV MCE protein-encoding genes are associated with two PqiA domains upstream. In the case of *E. coli* the predicted operon structures were *pqiA-pqiB-ymbA* and *yebS-yebT* (Fig. 3B), which resemble the most common type III and type IV MCE neighbourhoods, respectively. In these cases *ymbA* encodes the DUF330 domain containing protein and *pqiA* and *yebS* encode proteins with two PqiA domains. Each PqiA domain is predicted to span the inner membrane four times with N- and C-termini located in the cytoplasm^28,29^. PqiA is similar to NADH dehydrogenase subunit 2^30^, an antiporter domain involved in bidirectional membrane transport that requires energy from ATP hydrolysis but does not directly bind it. These data suggest that the mechanism of action of type III and IV transport complexes might differ from each other and type I and II proteins, an observation that is consistent with our previous structural data^14^.

### MCE proteins are integral inner membrane proteins

In agreement with our structural predictions for PqiB and YebT as envelope-spanning complexes, a recent study revealed that the bulk of PqiB and YebT are localised in the periplasm^31^. For both YebT and PqiB, the presence of a single transmembrane α-helix and the lack of a predicted signal sequence suggest the proteins are associated with the inner membrane (Supplementary fig. S3). This would be consistent with the demonstrated localisation of the type I MCE protein, MlaD, to the inner membrane^12^. However, YebT is the orthologue of the *V. parahaemolyticus* multivalent adhesion molecule, MAM-7, which was reported to be located in the outer membrane^20^. Therefore, it was essential to determine the cellular locations of PqiB and YebT. To probe their localization, inner and outer membranes of the parent strain, *E. coli* K-12 BW25113, and mutants lacking *pqiAB, yebST*, or both *pqiAB* and *yebST*, were fractionated and separated on sucrose density gradients. Using known outer and inner membrane markers, TolC and AcrB respectively, we demonstrated that PqiB and YebT could only be detected in the inner membrane fraction but not in either the outer membrane fraction or the corresponding mutants (Fig. 4). Therefore, we concluded that in *E. coli*, all MCE proteins are inner membrane associated proteins.

**Figure 4.**
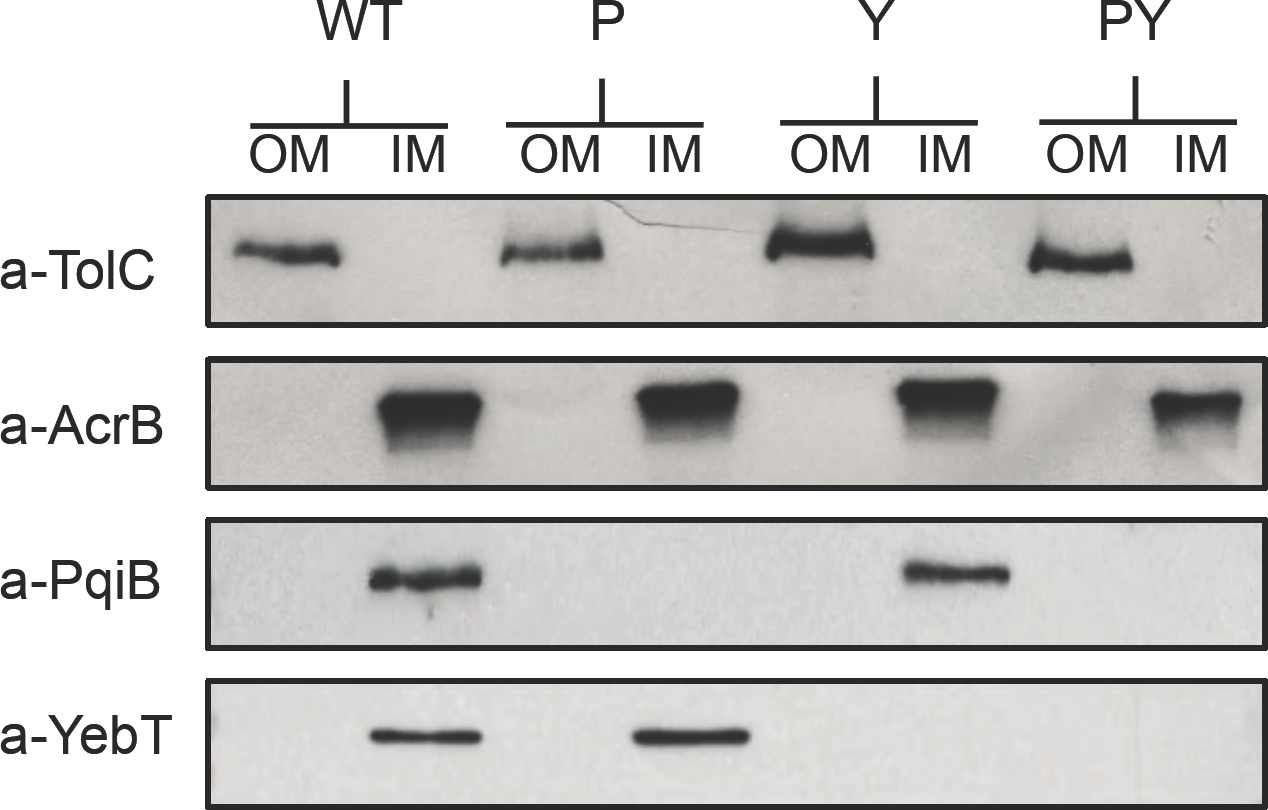
Western blots to identify the locations of PqiB and YebT in *E. coli* K-12 BW25113. The inner and outer membranes of the parental strain (WT) and the *pqiAB* (P), *yebST* (Y) and *pqiAB yebST* (PY) mutants were separated by sucrose gradients. The inner and outer membranes of each strain were taken as single fractions and separated by SDS-PAGE. Western blots using antibodies against known outer membrane marker TolC and inner membrane marker AcrB revealed that the separation of the inner and outer membranes was successful. Anti-PqiB and anti-YebT antibodies were used to identify the locations of PqiB and YebT.

### Loss of MCE proteins disturbs cell envelope homeostasis

Given the homology between the MCE proteins, we hypothesized that type III and type IV MCE proteins would contribute to cell envelope homeostasis in a manner consistent with the type I MCE protein, MlaD. Loss of outer membrane homeostasis in an *E. coli mlaD* mutant is indicated by the inability of the mutant to grow in the presence of SDS-EDTA^12^. However, in contrast the *E. coli pqiAB* and *yebST* mutants were as resistant to SDS-EDTA as the parent strain (Supplementary fig. S4). In an attempt to understand more about the roles of PqiB and YebT, we compared the growth of the parent strain and isogenic *pqiAB, yebST* and *pqiAB yebST* mutants in over 1900 growth conditions using BiOLOG Phenotype Microarrays. Phenotypes were identified for five compounds: lauryl sulfobetaine (LSB), tetracycline, penimepicycline, azlocillin and clioquinol (Fig 5A). However, in subsequent growth experiments, clear phenotypic differences were confirmed only for LSB. Growth of both the *pqiAB* and *pqiAB yebST* mutants was inhibited by 1% LSB, but the *yebST* mutant and the BW25113 parent strain were unaffected (Fig. 5).

**Figure 5.**
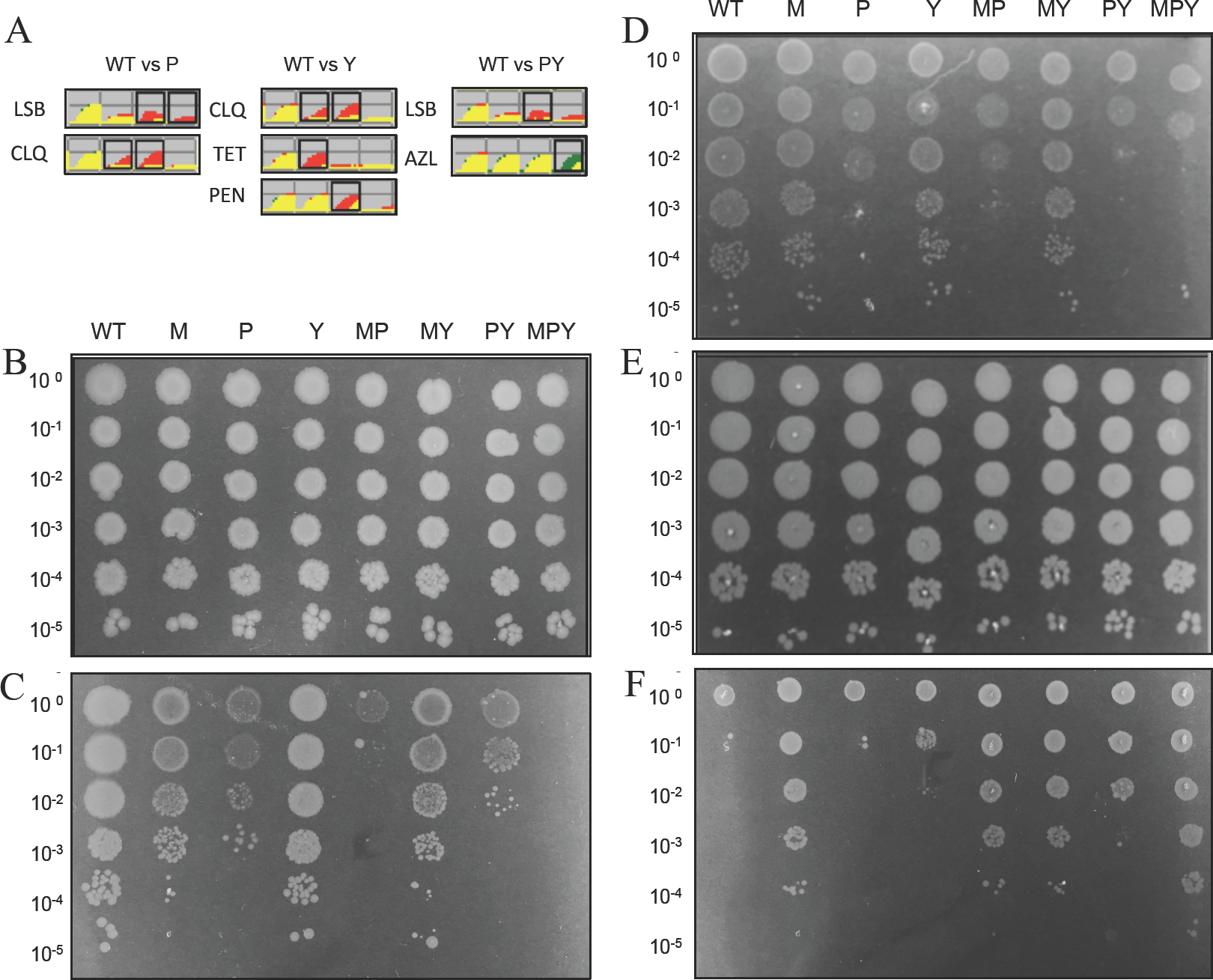
The phenotypes of MCE mutants on sulfobetaines and vancomycin. A) Selected results from the BiOLOG analysis: Red = growth of the parent strain, green = growth of the mutant, LSB=lauryl sulfobetaine, CLQ=clioquinol, TET=tetracycline, PEN=penimepicycline, AZL=azlocillin. B-F) Logarithmic dilutions of the parent and mutant strains on (B) LB agar, (C) 1% LSB, (D) 1% caprylyl sulfobetaine, (E) 1% myristyl sulfobetaine and (F) 300 µg/ml vancomycin. WT= parent strain, M= Δ*mlaD*, P= Δ*pqiAB*, Y= Δ*yebST*, MP= Δ*mlaD* Δ*pqiAB*, MY= Δ*mlaD* Δ*yebST*, PY= Δ*pqiAB* Δ*yebST* and MPY= Δ*mlaD* Δ*pqiAB* Δ*yebST*

A complete set of double mutants and the *mlaD pqiAB yebST* triple mutant was then constructed and screened for growth inhibition by 1% LSB (Fig. 5C). Like the *pqiAB* mutant, the single *mlaD* mutant was also sensitive to LSB. In addition, the *mlaD pqiAB* double mutant was more sensitive than either single mutant, revealing additive phenotypes for these strains. The *yebST* mutant was not sensitive to LSB, but this mutation further increased LSB sensitivity in the *pqiAB* strain, suggesting that YebST plays a minor role in LSB resistance. Consistent with this, we observed no growth at all for the triple mutant, the only strain to completely lack MCE domains. Complementation of these mutants restored growth on LSB, demonstrating that the absence of MCE proteins was the cause of this sensitivity (Supplementary fig. S5). These results show that all MCE proteins contribute to LSB resistance.

These data clearly show distinct roles for the *E. coli* MCE proteins in maintaining cell envelope homeostasis. We hypothesised that these differences may be due to differences in substrate specificity, such as variations in fatty acid chain length. To test this hypothesis we screened the mutants for growth on sulfobetaines with varying carbon chain lengths; caprylyl sulfobetaine and myristyl sulfobetaine (Fig. 5D and E; supplementary fig. S6). None of the strains were sensitive to myristyl sulfobetaine and only the *pqiAB* mutant was sensitive to caprylyl sulfobetaine. Similar to the screens on LSB, the *pqiAB yebST* double mutant and the triple mutant were more sensitive than the *pqiAB* mutant, revealing another additive phenotype for this strain. However, additive phenotypes were not observed for the other double mutants.

Sensitivity to detergents can often indicate loss of outer membrane integrity. To screen for such outer membrane defects, strains lacking one or a combination of MCE domain proteins were assayed for sensitivity against vancomycin, which does not typically cross the outer membrane. Surprisingly, our data revealed a large increase in vancomycin resistance for all *mlaD* mutants compared to the parent strain (Fig. 5F). This was also observed in the *pqiAB yebST* double mutant. Furthermore, the triple *mlaD pqiAB yebST* mutant was marginally more resistant than the other *mlaD* mutants. From these data, we conclude that the MCE proteins in *E. coli* have distinct but overlapping functions.

### MCE proteins contribute to maintenance of lipid asymmetry

MlaD was previously demonstrated to have a role in maintaining the lipid asymmetry of the outer membrane^12^. Given the overlapping functions of MlaD, PqiB and YebT, we hypothesized that PqiB and YebT may also be involved in maintaining outer membrane lipid asymmetry. To test this hypothesis we used the activity of the enzyme PagP as an indirect measure of surface exposed phospholipids; the enzyme converts lipid A from the hexa- to hepta-acylated form only when phospholipids are located in the outer leaflet of the outer membrane^32^. Previously it was demonstrated that in a Δ*mla* background, loss of lipid asymmetry could be enhanced by loss of the outer membrane phospholipase PldA^12^. Therefore, radiolabelled lipid A was isolated from the *mlaD, pqiAB*, and/or *yebST* mutants in otherwise wild-type and *pldA* backgrounds and separated by thin layer chromatography. As a positive control, the parent strain was treated with the chelating agent EDTA, which is known to result in increased hepta-acylation of lipid A^33^. As previously reported^12^, there was an increase in hepta-acylated lipid A relative to the parent in all strains lacking MlaD, and this effect was elevated in the absence of *pldA* (Fig. 6). However, the amounts of hepta-acylated lipid A in the *pqiAB, yebST* or *pqiAB yebST* mutants did not change when compared to the parent strain, suggesting that both PqiAB and YebST do not play major roles in maintaining outer membrane lipid asymmetry. Similarly, there was no increase in hepta-acylated lipid A in the *mlaD pqiAB* and *mlaD yebST* double mutants relative to the *mlaD* single mutant. These results are consistent with the fact that only the *mlaD* strain, but not the *pqiAB* or *yebST* mutants, is sensitive to SDS-EDTA (Supplementary fig. S4). We did, however, observe that there was a significant increase in the levels of hepta-acylated lipid A in the *mlaD pqiAB yebST* triple mutant when compared with the *mlaD* single mutant, particularly in the *pldA* background. These results indicate that cells lacking all MCE domain proteins show a larger asymmetry defect than those lacking just MlaD. We conclude that under standard laboratory conditions PqiB and YebT may contribute to outer leaflet integrity in the absence of the Mla pathway, but that their primary roles are clearly distinct.

**Figure 6:**
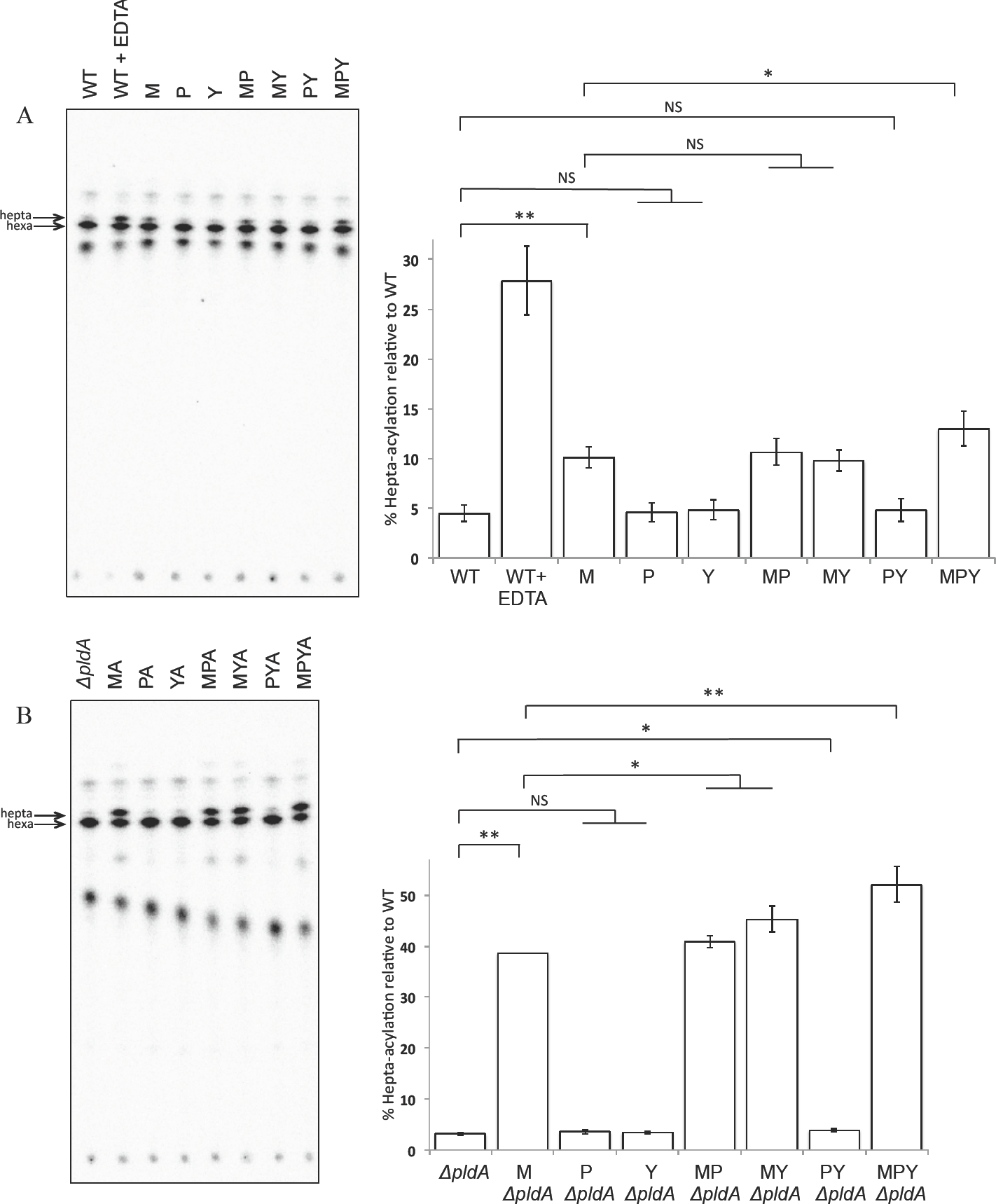
Hepta-acylation of lipid A in all MCE mutants and the parent strain as a measure of phospholipid accumulation on the cell surface. A) Lipid A analysis of MCE mutants in an otherwise WT background. B) Lipid A analysis of the MCE mutants in a *pldA* mutant background. Radiolabelled lipid A was extracted from the strains and separated by TLC (solvent system: 50:50:14.6:4.6 chloroform: pyridine: 96% formic acid: water). The bar chart represents the data from three biological replicates and the errors bars represent the standard deviation. Statistical significance was determined using a t-test and corrected for multiple testing, where NS=non-significant, q>0.05, *= significant, q<0.05 and **=significant, q<0.01. The q values from pairwise comparisons were as follows: WT vs M = 0.0000051, WT vs P = 0.45, WT vs Y= 0.35, M vs MP = 0.32, M vs MY = 0.35, WT vs PY = 0.35, M vs MPY = 0.011, *ΔpldA* vs M *ΔpldA* = 0.0000001, *ΔpldA* vs P *ΔpldA* = 0.23, *ΔpldA* vs Y*ΔpldA* = 0.26, M*ΔpldA* vs MP*ΔpldA* = 0.035, M*ΔpldA* vs MY*ΔpldA* = 0.014 and M*ΔpldA* vs MPY*ΔpldA* = 0.006.

## Discussion

Here, for the first time, we have provided in depth bioinformatic analyses of MCE proteins. We have demonstrated that these proteins are widely distributed across diderm bacteria, that they were present early in such bacterial species^34^ and evolved vertically with no major horizontal gene transfer events. Previous work suggested that MCE proteins in pathogenic Gram-negative bacteria possessed either six or seven MCE domains^20^. In contrast, we reveal the majority of Gram-negative bacteria possess MCE proteins with one domain and multi MCE domain-containing proteins, usually with 3 or 7 domains, are specific to Proteobacteria having evolved from a protein with a single MCE domain; other variants are much less common. We have identified the most common MCE protein architectures and demonstrated that these are associated with membrane transport functions.

We demonstrated that the *E. coli* MCE proteins, and by implication all MCE proteins, are associated with the inner membrane of diderm bacteria. This result is consistent with the lack of signal peptides in these proteins (Supplementary fig. S3)^35^. Our findings differ from previously published work on the *V. parahaemolyticus* YebT homolog (MAM-7), which is reported to be a surface-localised integral outer membrane protein^20^. Whilst we do not have an explanation for this discrepancy, our results are in agreement with other observations, including known features of inner membrane associated proteins^36^, our structural data^14^, and the published locations of MlaD^12,17^, PqiB^31^, YebT^31^ and the MCE proteins in the evolutionary divergent chloroplast^18^.

The common location of the MCE proteins, and the homology between them, suggested they possess similar functions. Here we show additive defects for growth on detergents. While sensitivity to detergents can indicate a general outer membrane permeability defect, the lack of sensitivity to the common detergent SDS (Supplementary fig. S7), and the resistance to vancomycin suggests this is not the case for bacteria deficient in MCE proteins. Furthermore, the phenotypes observed here for the growth of the mutants on sulfobetaines with different carbon chain lengths suggest MCE proteins have an overlap in function but they also highlight mechanistic differences. Indeed, LSB is known to inhibit the carnitine/acyl-carnitine transporter in mitochondria as a substrate analog^37^. A potential explanation for our observed phenotype is that LSB inhibits a similar lipid transport pathway in *E. coli*, perhaps one that has some overlap in function to the MCE pathways. Alternatively, MCE proteins might be involved in trafficking LSB away from its target; therefore the variable phenotypes observed on alternative sulfobetaines would represent differences in substrate specificity. Indeed, a type I operon in *Sphingobium japonicum* is essential for the uptake and utilisation of *γ*-hexachlorocyclohexane^15^ and a type I operon in *Pseudomonas putida* is associated with toluene resistance^16^, suggesting there is a large range of substrates for MCE proteins, not simply membrane lipids.

While the exact functions of PqiB and YebT are yet to be determined, our recent data reveals that they share the same lipid-binding properties as MlaD^14^. Thin layer chromatography (TLC) and mass spectrometry data revealed phosphatidylglycerol and phosphatidylethanolamine bound to all three MCE-domain containing proteins. Cryo-EM structures revealed that all three proteins form homo-hexamers with a hydrophobic central pore that could harbour phospholipids or other hydrophobic molecules^14^. Furthermore, our recent observation that the cell surface of the triple mutant is visibly ruffled^14^, which has also been observed in *Streptomyces*^38^ and *Mycobacteria*^8^, indicates the phenotypes observed for the triple mutant are a consequence of the increasing perturbation of the cell envelope. Indeed, the SDS-EDTA sensitivity of the *mlaD* mutant^12,31^, and the exacerbation of this phenotype in the absence of the other MCE proteins^31^, suggests that the MCE proteins are involved in maintaining outer membrane homeostasis.

However, the precise contribution of the different MCE proteins to the outer membrane integrity is not understood. Despite their similar affinities for phospholipids there are no appreciable differences in the ratios of the major phospholipids in the outer membrane of the triple mutant when compared to the WT (Supplementary fig. S8). Nevertheless, it remains possible that there are differences in the total amount of phospholipid in the outer membrane, a hypothesis that is yet to be tested. However, previous studies demonstrated that an *mlaD* mutant accumulates phospholipids on the cell surface^12^ supporting a role for MCE domain proteins in maintaining outer membrane asymmetry. Although we could recapitulate these observations, and demonstrate a further increase in phospholipids in the outer leaflet in a triple mutant, no appreciable differences were observed between the parent, *pqiAB* or *yebST* mutants. This suggests that the primary roles of PqiB and YebT differ to that of MlaD. Indeed, the MCE proteins may function in different environmental niches or conditions. We have only tested their function under standard laboratory growth but there is evidence that the three operons are differentially expressed^39,40^. Furthermore, it is well known that bacteria can modulate the lipid content of their cell envelope under different environmental parameters^41^. Thus, it remains possible that PqiB and YebT are important for maintenance of lipid asymmetry in conditions yet to be tested. Notwithstanding their enigmatic function, their importance as crucial components of diderm bacteria is supported by the fact these operons remain intact in organisms such as *Salmonella enterica* Typhi which are undergoing genome reduction due to their host restricted lifestyles^42^.

In conclusion, we have shown that MCE domains arose early in the evolution of diderm bacteria in the form of single-domain MCE proteins, later evolving as multi-domain MCE proteins in Proteobacteria. Through phenotypic studies, we have shown that the roles of multi-domain MCE proteins in Proteobacteria overlap with the role of single-domain MCE proteins, but that these proteins also fulfil other roles within the bacterial cell. Based on the data presented here, we propose that multidomain MCE proteins fulfil a role as components of a novel type of phospholipid transporter, homologous to both the MCE ABC transporters and antiporters, and are involved in the maintenance of the Gram-negative cell envelope.

## Methods

### Architecture definition and phylogenetic distribution

MCE domain-containing sequences were retrieved by scanning the UniProtKB database^19^ (version 2015_03) for matches to the PFAM MCE HMM^1^ (accession PF02470.15) using HMMER (hmmsearch 3.1b2)^21^. These thresholds were judged appropriate through manual inspection of known MCE domain-containing *E. coli* proteins. MCE protein sequences were re-scanned using HMMER to scan for all other PFAM domains (Pfam database 27.0), using the PFAM gathering bit score thresholds. Only the most significant hit was retained in overlapping predictions with members of the same clan. MCE domain hits were retained in all cases. These resulting predicted domains were sorted by start position and were used to define protein architectures.

To provide a percentage measure of prevalence for each MCE architecture in each taxonomic rank, species with fully sequenced genomes (according to UniProt) were examined for the absence or presence of each MCE architecture. To minimise bias due to the sequencing of many strains in well-studied species such as *E. coli*, one genome sequence per species was selected at random. These results were then summed for each species phylum to calculate the percentages.

### Clustering

A representative set of 1,734 MCE proteins selected using the CD-HIT Suite at a cut-off of 50% protein identity were used in all-against-all searches in BLASTP with an e-value threshold of 1e-15. Information about each protein (phylum, architecture, etc) was incorporated into the BLAST results based on the previous architecture designations and information from UniProt. Cluster diagrams were constructed using Cytoscape (v.3.3.0), where each node represents a single protein sequence and each line (or edge) represents a match below the e-value threshold.

### Gene neighbourhoods

The Ensembl gene ID for each MCE protein in Proteobacteria was cross-referenced in conjunction with the NCBI taxonomic identifier to retrieve the sequence database for each organism in Ensembl (using the Ensembl Perl API)^43,44^. Information was retrieved for up to 10 genes located up to 10 kb eitherside of the MCE gene. Domain architectures were predicted for the encoded proteins by scanning for PFAM domains (PFAM database 27.0) using HMMER (hmmsearch 3.1b2). Where several genomes for a particular species were found, a single representative genome was chosen randomly so that each species was only represented once in the neighbourhood results.

To display groups of conserved neighbourhoods together, gene neighbourhoods were first clustered using a nearest neighbour joining approach. The similarity measure used in the clustering method gives greater weight to genes closer to the centre of the neighbourhood, based on the assumption that gene positions further away from the centre of the neighbourhood are less likely to be conserved.

This clustering method was used to constructs neighbourhood diagrams for each protein architecture, where similar neighbourhoods were clustered together. The most common neighbourhoods were manually selected from these diagrams, and some gene domain architectures were combined (e.g. “ABC_tran” domain architecture was merged with “ABC_tran, AAA_21” architecture due to similar predicted functions and similar neighbourhoods of these genes). The neighbourhoods were split into type I, III and IV proteins. To calculate a percentage for a particular gene neighbourhood, the number of genes with that neighbourhood was divided by the total number of genes that encode the given protein architecture.

### Operon predictions

Operons of *pqiB* and *yebT* were predicted using ProOpDB^45^ and EcoCyc^46^. The domain architecture for each protein encoded in these operons was predicted using HMMER.

### Strains, media and growth conditions

*E. coli* K-12 BW25113 was used as the parent strain^47^. Bacteria were grown in Luria-Bertani (LB) medium or on LB plates (LB supplemented with 1.5% nutrient agar) and incubated at 37°C. If required, the medium was supplemented with 50 µg/ml kanamycin or 100 µg/ml carbenicillin. To construct deletions in *pqiAB* and *yebST*, the genes were replaced by a kanamycin resistance cassette, as previously described^48^. To construct gene deletions in *mlaD,*the *mlaD::aph* was transferred from the Keio collection^47^ by P1 transduction as described previously^49^. The kanamycin cassette was removed using the vector pCP20^48^. For dilution plates, overnight cultures were adjusted to an OD_600_ of 1 and diluted down to 10^−5^ in a microtitre plate. A multichannel pipette was used to transfer 2 µl of the dilutions for each strain to LB agar plates supplemented with the desired chemicals.

### Separation of the inner and outer membranes for identification of cellular location

This method was adapted from the methods described by Osborn & Munsen, 1974^50^ and Daleberoux et al., 2015^51^. For each strain required, 2 litres of cells were grown to an OD_600_ of 0.6-0.8. The cells were pelleted (16,000 *g* for 10 minutes) and re-suspended in 10 ml of sucrose-Tris buffer (0.75 M sucrose, 10 mM Tris, pH 7.8). To form spheroplasts, the mixture was transferred to an Erlenmeyer flask where 500 µl of 2 mg/ml lysozyme was added. After 2 minutes of incubation on ice, 20 ml of ice-cold 1.5 mM EDTA was slowly added over 10 minutes using a peristaltic pump, with gentle stirring. The spheroplasts were broken using a C3 cell disrupter (at 15000 Psi) and unbroken cells were pelleted at 17,400 *g* for 20 minutes. The supernatant was spun again at 48,400 g for 1 hour to pellet cell membranes. To wash the membranes, the pellet was re-suspended in 20 ml of sucrose-Tris-EDTA buffer 1 (0.25 M sucrose, 3.3 mM Tris, pH 7.8, 1 mM EDTA) and re-pelleted at 165,000 *g* for 1 hour. The membranes were re-suspended again in 10 ml of sucrose-Tris-EDTA buffer 2 (20% sucrose, 0.5 mM EDTA, 10 mM Tris, pH 7.8).

For the gradient, all sucrose was dissolved in EDTA-Tris buffer (0.5 mM EDTA, 10 mM Tris, pH 7.8). The gradient was made up in 38.5 ml Ultra-Clear Thinwall tubes (Beckman Coulter) with 10 ml of 73% sucrose (bottom), 18 ml of 53% sucrose (middle) and the 10 ml of membrane sample in 20% sucrose (top). The gradient was centrifuged at 141,000 *g* for 40 hours in a SW 28 Ti rotor at 4°C. To obtain the inner membrane a pipette was used to withdraw the membrane through the top of the gradient from the 20%-53% boundary. To obtain the outer membrane the tube was pierced at the bottom and the membrane was collected by gravity flow from the 53%-73% boundary.

The resulting isolated fractions were analysed by western blotting following protein separation by SDS-PAGE. Antibodies against the known membrane markers TolC (outer) and AcrB (inner) were used to assess the success of separation. Antibodies against PqiB and YebT were used to determine the locations of PqiB and YebT. All primary antibodies were used at a dilution of 1:2,000 in Tris-buffered saline and left to incubate overnight. After washing, the blots were labelled with horse radish peroxidase secondary antibody (Sigma Aldrich) (dilution 1:15,000). The western blots were developed using ECL Prime Western Blotting Detection Reagent (Amersham), and exposed to Hyperfilm ECL (Amersham) for between 5 seconds and 5 minutes.

### Analysis of lipid A

Lipid A was extracted and analysed as described previously^52^. In brief, 5 ml cultures were grown to OD_600_ 0.6-0.8 with 1 µCi/ml [32P]-disodium phosphate and harvested by centrifugation. One parent culture was treated with 25 mM EDTA, pH 8.0 for 10 minutes before harvesting. The pellet was washed twice with PBS and finally converted into a single phase Bligh-Dyer^53^ mixture (1:2:0.8 chloroform:methanol:water). After a 20-minute incubation at room temperature, the mixture was centrifuged and the pellet washed once with 1 ml single phase Bligh-Dyer mixture. After another centrifugation the pellet was re-suspended in 12.5 mM sodium acetate containing 1% SDS and sonicated for 15 minutes before incubation at 100°C for 40 minutes. The mixture was then converted into a two-phase Bligh-Dyer mixture (2:2:1.8 chloroform:methanol:water). After centrifugation, the lower phase of each mixture was collected and washed with 1 ml of fresh upper phase prepared from a two-phase Bligh-Dyer mixture. The final lower phase was collected after centrifugation and dried under nitrogen gas. The dried samples were re-dissolved in 100 µ1 of 4:1 chloroform:methanol and 20 µ1 of the sample was used for scintillation counting. Equal amounts of radiolabeled lipids were spotted onto a silica TLC plate and were separated using 50:50:14.6:4.6 chloroform:pyridine:96% formic acid:water. The TLC plate was dried and exposed to phosphor storage screens (GE Healthcare) and was visualised in a phosphor-imager (Storm 860, GE Healthcare). The spots were analyzed by ImageQuant TL analysis software (version 7.0, GE Healthcare). Spots were quantified and averaged based on three independent experiments of lipid A isolation. Before performing statistical tests the datasets were first tested for normality using a Shapiro-Wilk test. A one-sided unpaired t-test was then used to determine statistical significance between pairs of samples. To correct for multiple testing the Benjamini-Hochberg correction was applied to all p values.

### Lipid extraction and thin layer chromatography

Lipids were extracted by Bligh-Dyer^48^ from outer membranes fractions prepared by sucrose gradient (as described above). A total of 5.7 ml of 1:2 chloroform: methanol was added to the whole outer membrane fraction (approx. 1.5 ml), followed by 1.875 ml of chloroform, followed by 1.875 ml water, with thorough mixing at each stage. The mixture was centrifuged at 1000 rpm in an IEC table-top centrifuge for 5 minutes and the lower (organic) phase was collected using a glass Pasteur pipette and transferred to a new tube. To prepare fresh upper phase, the same procedure was repeated but with 1.5 ml of water (instead of sample). The upper phase was collected and mixed at a 2.25:1 ratio with the already obtained lower phase (extracted from the outer membrane). After further centrifugation, the lower phase was collected and transferred to a fresh tube. The liquid was dried under nitrogen gas and the lipids were re-dissolved in 200 µl of chloroform. For thin layer chromatography, 10 µl of extracted lipid was pipetted onto 7 x 7 cm silica gel 60 plates and separated by two different solvent systems (direction 1=65:25:4 chloroform: methanol: water, direction 2=80:12:15:4 chloroform: methanol: acetic acid: water). After thorough drying the plate was sprayed with phosphomolybdic acid and heated using a heat gun until lipids were clearly visualised.

### Data availability

The BiOLOG datasets generated during this study are available from the corresponding author on reasonable request.

## Acknowledgments

This work was supported by a BBSRC grant to IRH. GLI was funded by the MIBTP BBSRC PhD Scholarship. This work was supported in part by the Singapore Ministry of Education Academic Research Fund Tier 2 (MOE2013-T2-1-148) grant (to SSC). The authors thank Dr. Alan McNally for critical reading of the manuscript.

## Author contributions

GLI conducted the experimental work with help from ZSC, JAB, MJ, TJK and MS. NJD did all of the bioinformatics. GLI and IRH wrote the manuscript with contributions from NJD, ZSC, AFC, SSC and JAC. GLI lead the project with supervision from JAC and AFC, and IRH who conceived the study. All authors reviewed the manuscript.

## Additional information

**Competing financial interests:** The authors declare no competing financial interests.

